# Blue light advances bud burst in branches of three temperate deciduous tree species under short-day conditions

**DOI:** 10.1101/259101

**Authors:** Craig C. Brelsford, T Matthew Robson

## Abstract

During spring, utilising multiple cues allow temperate tree species to coordinate their bud burst and leaf out, at the right moment to capitalise on favourable conditions for photosynthesis. Whilst the effect of blue light (400-500nm) has been shown to increase percentage bud burst of axillary shoots of *Rosa* sp, the effects of blue light on spring-time bud burst of temperate deciduous tree species has not previously been reported. We tested the hypotheses that blue light would advance spring bud burst in temperate tree species, and that late-successional species would respond more than early-successional species, who’s bud burst is primarily determined by temperature. The bud development of *Alnus glutinosa, Betula pendula*, and *Quercus robur* branches, cut from dormant trees, was monitored under two light treatments of equal photosynthetically active radiation (PAR, 400-700 nm) and temperature, either with or without blue light, under controlled environmental conditions. In the presence of blue light, the mean time required to reach 50% bud burst was reduced by 3.3 days in *Betula pendula,* 6 days in *Alnus glutinosa,* and 6.3 days in *Quercus robur*. This result highlights the potential of the blue region of the solar spectrum to be used as an extra cue that could help plants to regulate their spring phenology, alongside photoperiod and temperature. Understanding how plants combine photoreceptor-mediated cues with other environmental cues such as temperature to control phenology is essential if we are to accurately predict how tree species might respond to climate change.

**Key Message:** An LED spectrum containing blue light advanced bud burst in branches of *Betula pendula, Alnus glutinosa* and *Quercus robur* compared with a spectrum without blue light in a controlled environment.

## Introduction

Trees at northern latitudes must time the bud burst and unfolding of their leaves in spring so that they are able to capitalise on favourable conditions for photosynthesis as early as possible, whilst avoiding late frosts which can cause foliar damage (Hänninen 1991, Augspurger 2009, Bennie et al. 2010). Prior to bud-burst, a tree must go through three dormancy periods over winter which can be separated into 1) paradormancy – the inhibition of growth by distal organs, 2) endodormancy - the inhibition of growth by internal bud signals, and 3) ecodormancy - the inhibition of growth based on temporarily-unfavourable light conditions (Samish 1954, Lang et al. 1987,). The dormancy period between endodormancy and ecodormancy is jointly controlled by a chilling requirement and by photoperiod, where the chilling requirement is defined as non-freezing temperatures below 10°C (Battey 2000). After or concomitantly with this period of vernalisation, temperature is the predominant cue affecting bud burst, and to predict bud burst many models calculate the accumulated daily mean temperature above a given minimum threshold (e.g. >0°C) after sufficient vernalisation has occurred (Hänninen 1995, Cesaraccio et al. 2004).

Typically, early-successional light-demanding species adopt a risky strategy by timing their bud burst primarily based on temperature (Körner 2007, Caffarra and Donnelly 2010, Körner and Basler 2010), whereas it is more common for later-successional shade-tolerant species to be photoperiod sensitive (Basler and Körner 2012). Körner and Basler (2010) propose that in a warmer climate the absence of a photoperiod requirement in opportunistic early-successional species will probably give them a competitive advantage over later-successional species. However, this may also lead early-successional species to be at greater risk from late spring frosts (Cannell and Smith 1986), since a day-length threshold provides a fail-safe mechanism ensuring that a period of unseasonably warm weather during the winter cannot prematurely fulfil the temperature requirement. The date of bud burst in tree species is considered to have advanced by 2.5 days per decade at temperate latitudes since 1971 due to climate change (Menzel et al. 2006, Körner and Basler, 2010), however this trend for promotion of bud burst date with warmer temperatures has recently started to diminish (Fu et al. 2015). Specifically, leaf unfolding was advanced by fewer days per additional °C of warming in 13 temperature deciduous woody species measured across northern temperate Europe and the USA from 1980-2012. Fu et al. (2015) partially attributed this decline to fewer chilling days, but also to a photoperiod requirement that reduces the influence of warmer springs. Additionally, periods of “global dimming” caused by anthropogenic and volcanic emissions, affect the spectral composition of incident solar radiation reaching the Earth’s surface (Stanhill and Cohen 2001, Mercado et al. 2009), increasing its diffuse/direct fraction and consequently also the relative proportion of blue light (420-490 nm) (Urban et al. 2007). If we are to forecast the response of bud burst to climate change, we will need to improve existing models to include and account for all the mechanisms involved (Way and Montgomery 2014, Yun et al. 2017, Mayshev et al. 2018).

The effect that spectral composition has on the timing of bud burst and leaf out in tree species has been studied much less than that of photoperiod. The spectral composition that trees receive changes predictably with time of year, day length, and latitude; and over shorter time-scales with cloud cover and other factors that alter the ratio of direct to diffuse solar radiation (Johnson et al. 1967, Smith 1982, Hughes et al. 1984, Urban et al. 2007). These compositional changes have implications for tree species undergoing range shifts under rising global temperatures where differences in spectral irradiance at higher latitudes will produce novel combinations of phenological cues (Wareing 1956, Saikkonen et al. 2012). Recently, light pollution from street lamps has also been shown to advance the bud burst date of tree species *Acer pseudoplatanus*, *Fagus sylvatica, Fraxinus excelsior* and *Quercus robur* across the whole UK (Somer-Yeates et al. 2016). Although the regions of the spectrum produced by street lamps that were responsible for the earlier bud burst were not specified, this study highlights the potential ecological role for light quality influencing bud burst. What little research has questioned which regions of spectral irradiance are likely to provide cues for bud phenology has largely focussed on the ratio of red (R, 655-665 nm) to far-red light (FR, 725-735 nm, Smith 1982). For instance, FR light is enriched during the periods of twilight from dawn to sunrise and between sunset and dusk, and thus the R:FR ratio varies with day length, latitude and the time of year (Holmes and Smith 1975, Smith 1982, Hughes et al. 1984). In a controlled-environment study, simulating twilight with a treatment of FR light caused bud burst to advance in *B.pendula* (Linkosalo et al. 2006), implicating phytochromes in the coordination this response. However, bud burst mediated by R and FR light has been shown to be both species-specific and dependent on the population studied (Erez et al. 1966, Mølmann et al. 2006, Girault et al. 2008).

Numerous studies have investigated how R and FR light affect bud burst and yet the mechanisms behind this response are not well understood, far less still is known about the effects of blue light on bud burst. The blue light region of solar radiation is also enriched during the hours before sunrise and after sunset (Robertson 1966, Johnson et al. 1967, Hughes et al. 1984). Since cryptochrome photoreceptors, that detect blue and ultraviolet-A radiation (UV-A, 315-400 nm), are known to influence the circadian rhythm and photoperiodic flowering responses (Guo et al. 1998, Somers et al. 1998). This result implies that it could be worthwhile testing whether blue light can serve as a cue for bud burst in temperate deciduous tree species. Of the few studies on the effects of blue light on bud burst, Girault et al. (2008) report that blue light (Philips TL-D 18W/18 fluorescent tubes) was as effective as white light (Osram L 18W/77 Fluora fluorescent tubes, 31% blue; both spectra given in SI of Girault et al. 2008) in terms of the percentage of axillary buds reaching bud burst in *Rosa* sp. However, the timing of spring bud burst in trees may not be mediated through the same mechanisms as axillary buds of herbaceous species: for instance, monochromatic blue light (460nm) did not induce bud-burst in the gymnosperm *P.abies* after 6 weeks of 24h photoperiod (Mølmann et al. 2006). To our knowledge, there have been no studies investigating the effects of blue light on the bud burst of temperate deciduous tree species.

Given that blue light has been reported to increase the percentage bud burst in the vegetative shoots of angiosperms including *Rosa* sp. and another Rosaceae, *Prunus cerasifera* (Muleo et al. 2000, Girault et al. 2008), we hypothesised that blue light would advance bud burst in temperate deciduous tree species. Additionally, we expected that bud burst of early successional species, which is predominantly affected by temperature (Basler and Körner, 2012), would be less affected by blue light than late successional species. To this end, we tested the effect on bud burst of light treatments that either included or excluded blue light in branches of the three temperate deciduous species *A.glutinosa, B.pendula* and *Q.robur*, differing in their functional strategy from early successional to late successional.

## Methods

### The collection and care of branch material

We collected dormant branches of *A.glutinosa, B.pendula* and *Q.robur* from co-occurring populations of local origin in a natural forest stand in southern Finland (60°13’04.2“N 25°00’31.0“E) and kept them in a temperature-controlled environment under different light treatments while monitoring their bud development. Branches were collected from the lower canopy of trees (2-3 m height above the ground) using sterile garden secateurs on 6^th^ February 2017. We selected eight trees per species, taking 10-15 branches from each tree. Only dormant branches, free of pests and symptoms of disease were selected.

The use of this approach, putting cut twigs in controlled conditions for the study of bud burst phenology, and its reliability have recently been exemplified in several studies (Basler and Körner 2012, Laube et al. 2014, Polgar et al. 2014, Vitasse and Basler 2014, Primack et al. 2015). We followed a similar protocol to these studies, placing the cut end of twigs immediately into test tubes of water and continuously monitoring their bud development, under two light treatments: a broad spectrum containing blue light, and a broad spectrum of equal PAR and far red but with blue light attenuated (Table S1).

All trees received 111 chilling days prior to harvest: this was calculated according to Murray et al. (1989) as the number of days from 1^st^ of November with a mean temperature ≤5°C. All trees were on the east-facing side of a mixed-forest stand, and only branches that were exposed to sunshine for the majority of the day were selected (spectral composition provided in Table S2, monthly precipitation and temperature in Table S3). Branches were selected which presented at least five buds including the terminal bud. Once collected, branches were all immediately taken into a greenhouse kept at ambient outdoor temperature (0°C), and re-cut to a mean length of 28.35 cm (mean ± 0.34 SE), and then placed into upright sterilised clear-plastic test tubes (10.26cm long – 15ml volume) containing tap water (Table S4, following Basler and Körner 2012 and Polgar et al. 2014). These twigs were directly removed into a temperature-controlled room with six compartments measuring 97-×-57-×-57 cm. Branches from individual trees of each species were randomly assigned into each compartment, so that in each compartment there were 16 branches of *Q.robur*, 20 branches of *B.pendula* and 16 branches of *A.glutinosa*. Twigs were rotated inside the compartments every two days to avoid any bias coming from undetected microclimatic variation inside the compartments. Lights were kept at the same distance from the tips of branches in each compartment.

### Light treatments under controlled conditions

Three of the six compartments received a broad spectrum of irradiance from an array of LED lamps (2 × Valoya AP67, 400-750 nm, Table S1: mean PAR among compartments of 161 ± ± ≤ 10 µmol m^-2^ s^-1^, plus 28 µmol m^-2^ s^-1^ far red) including blue light (BL, 21.8 µmol m^-2^ s^-1^, 13.5% of total PAR), and in the other three compartments BL was attenuated (no BL)by wrapping film Rosco #313 Canary Yellow (Westlighting, Helsinki, Finland) around the LED lamps (3 × Valoya AP67), limiting the spectrum to a wavelength range 500-750nm (Table S1: PAR 156± ≤ 10 µmol m^-2^ s^-1^, plus 31 µmol m^-2^ s^-1^ far red). Spectral irradiance measurements were taken inside all the experimental compartments with an array spectroradiometer (Maya 2000Pro, Ocean Optics Inc., Dunedin, Florida, USA) to ensure equivalent PAR and R:FR ratios between the treatments (Table S1). Lights were kept on a 12 h light-dark cycle to simulate photoperiod conditions experienced at the beginning of spring in southern Finland (60°13’04.2“N 25°00’31.0”E), to ensure that photoperiod would not be a confounding factor in the experimental test of light quality (Rinne et al. 1994, Welling et al. 2004, Søgaard et al. 2008).

### Temperature measurements under controlled conditions

Day/night temperature was thermostatically adjusted to maintain 12.6/9.5 ± 0.05 °c. Temperatures were monitored with iButton sensors (Maxim Integrated, San Jose, United States) in each compartment, sampling every one hour for the entire duration of the experiment, with a resolution of 0.065°C. An infra-red thermal thermometer (Optris LS LT, Berlin, Germany) was used to measure the temperature on individual buds of branches inside the compartments, to ensure there was no difference in bud temperature between treatments. The temperature of buds measured with the infra-red thermometer had a strong positive correlation with mean daily air temperature measured with iButton sensors (*R*^2^=0.83, Fig. S1, Table S5),.

### Monitoring bud development

Bud burst was scored until buds reached stage 4 on a scale of 1-7, made by adapting the scales from Teissier du Crois et al. (1981) and Kramer et al. (2017) whereby, 1 = dormant bud, 2 = bud swelling, 3 = bud scales begin to split, green visible on the bud, 4 =bud split, with leaf tip protruding. The experiment lasted 57 days at which point all species in all compartments had reached at least 50% bud burst (stage 4). Degree-days were calculated as the accumulated mean daily temperature > 0°C after the date of December 31^st^ 2016, following methods by Heide (1993), including any degree-days accumulated by the branches from the dates January 1^st^ to February 6^th^ 2017, during the period prior to the experiment.

### Data analysis

All results are given as the means ± 1 SE of three replicates, whereby each compartment was treated as one replicate. We followed statistical methods from Guak et al. (1998), Heide (2003), Søgaard et al. (2008), and Olsen et al. (2014), in comparing the days and degree-days needed to reach 50% bud burst in the BL and no BL treatments for each species. The effect of BL on the bud burst of each species was analysed using an ANOVA (R version 3.2.2., R Core Team, 2017). Pearson’s correlation was used to test the relationship between bud temperature and recorded air temperature in each compartment. Figs. 1 and 2 were produced in Microsoft Excel (2013).

### Results and Discussion

In all three species, BL decreased the mean number of days until 50% bud burst (Fig. 1). The response to blue light was smallest in the early successional species *B.pendula*, which reached 50% bud burst only 3-days earlier in the BL treatment compared to the no BL treatment (F_1_,_4_ = 100, *p* < 0.001). The later successional *A.glutinosa* and *Q.robur* reached 50% bud burst 6 days (F_1,4_ = 16.20, *p* = 0.016) and 6.3 days (F_1,4_ = 90.25, *p* < 0.001) earlier respectively in the BL treatment compared to the no BL treatment. These results are consistent with other studies showing that early-successional species achieve bud burst sooner than late-successional species (Wesołowski and Rowiński 2006, Basler and Körner 2012).

**Fig. 1.**
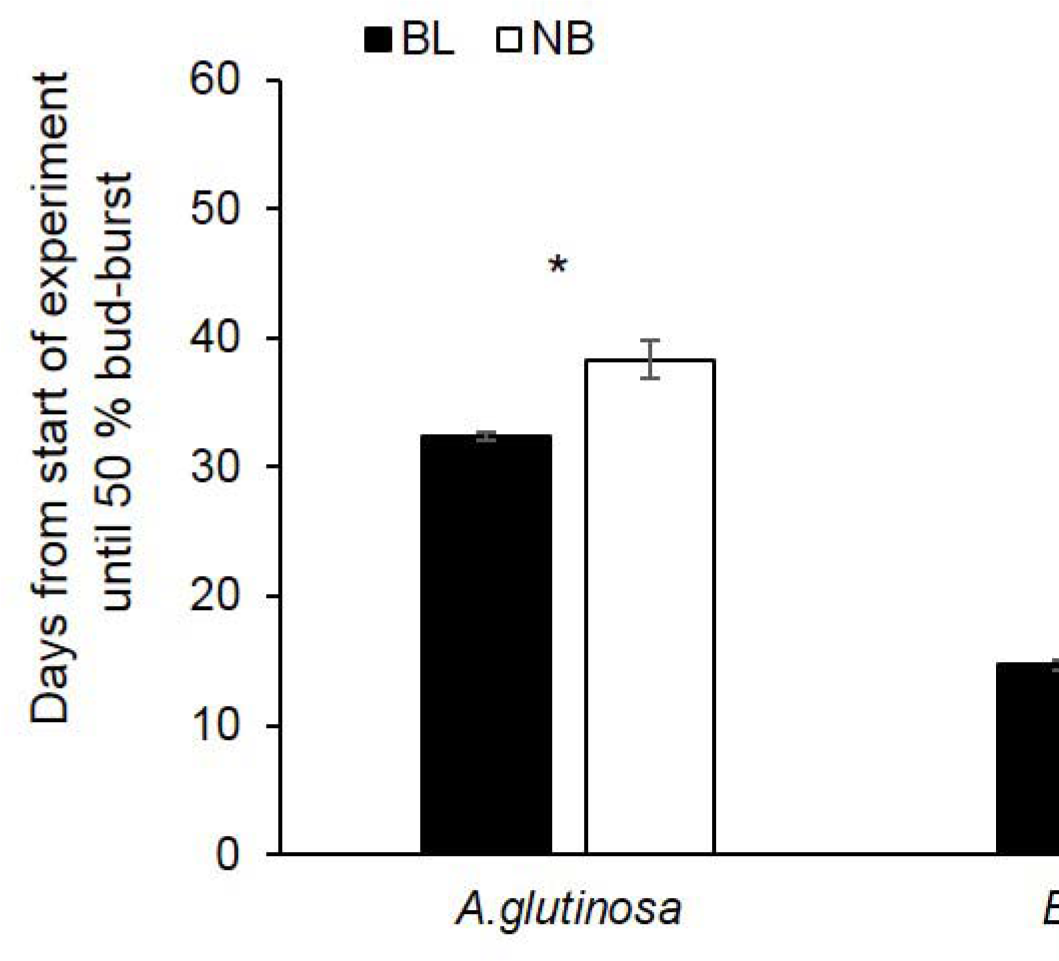
The mean number of days until 50% bud burst in *A.glutinosa, B.pendula* and *Q.robur*, in a broad spectrum treatment with blue light (BL) and without blue light (No BL). Error bars represent standard error (n=3) from three replicate compartments, each containing 16 *A.glutinosa* branches, 20 *B.pendula* branches and 16 *Q.robur* branches. Asterisks represent statistically significant differences between treatments, where * = *p* ≤ 0.05, ** = *p* ≤ 0.010, *** = *p* ≤ 0.001. *A.glutinosa* (F_1,4_ = 16.20, *p* = 0.016) *B.pendula* (F_1,4_ = 100, *p* < 0.001), *Q.robur* (F_1,4_ = 90.25, *p* < 0.001).

In our experiment, the effect of BL expressed as a degree day difference between treatments was consistent with the days until 50% bud burst (Fig. 2). Our BL light treatment reduced the degree-days required for bud burst in *B.pendula* by 17.12 degree-days (F_1,4_ =12.3, *p* = 0.025), in *A.glutinosa* by 28.14 degree-days (F_1,4_ = 35.8, p=0.04) and in *Q.robur* by 60.04 degree-days (F_1,4_ = 20.9, p= 0.010): this is the equivalent of a 10.54%, 15.48% and 11.03% reduction due to BL in each species respectively (Table 1). The degree-day requirement for bud burst in *B.pendula* was very similar to that of a natural population in southern Finland (Table S6), and was within the range reported in previous studies (Basler and Körner 2012, Heide 1993), as was *A.glutinosa* (Table S6, Heide 1993). The bud burst of *Q.robur* branches had a lower degree-day requirement than a natural population in southern Finland (Table S6): this discrepancy may derive from the photoperiod sensitivity of bud burst in *Quercus* species (Basler and Körner 2012, Basler and Körner 2014) which could have been activated by the 12h photoperiod we employed. Alternatively, this difference could be due to the reduced number of chilling days that treated branches received prior to February compared with those in forest stands throughout spring (Table S6). When expressed as the difference in the total length of time until bud burst, the effect of BL in days or degree-days was smallest in branches of *B.pendula:* however, because of the three species we compared it required the shortest total period before bud burst, the effect of BL on *B.pendula* was greatest in relative terms when measured as a % difference (Table 1).

**Fig. 2.**
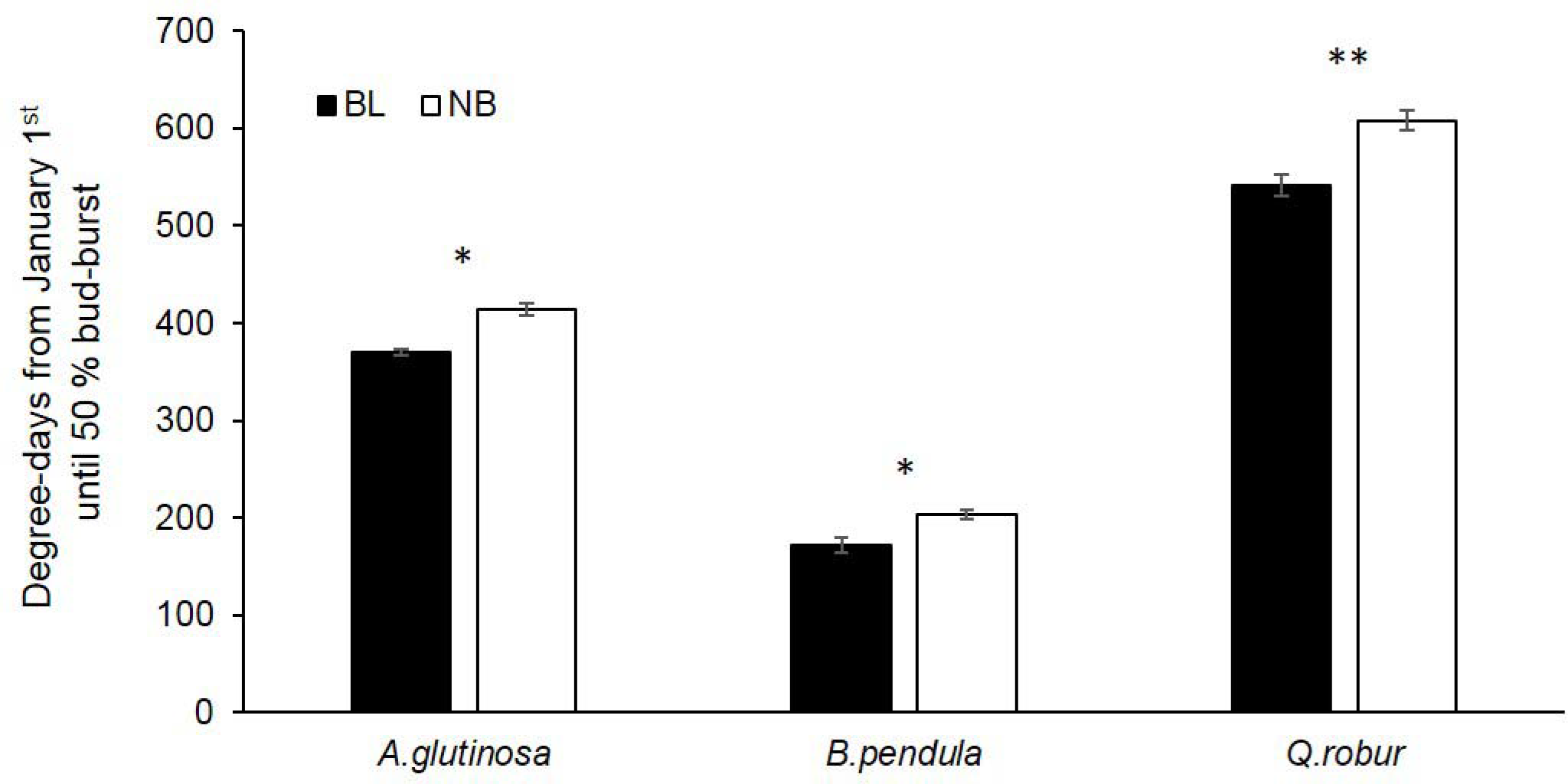
Mean degree-days until 50% bud burst in *A.glutinosa, B.pendula* and *Q.robur*, in a broad spectrum treatment with blue light (BL) and without blue light (No BL). Error bars represent standard error (n=3) from three replicate compartments, each containing 16 *A.glutinosa* branches, 20 *B.pendula* branches and 16 *Q.robur* branches. Degree-days calculated as the accumulated daily mean temperature < 0°C after the date of December 31^st^ 2016. Asterisks represent statistically significant differences between treatments, where * = *p* ≤ 0.05, ** = *p* ≤ 0.010, *** = *p* ≤ 0.001. *A.glutinosa* (F_1,4_= 35.8, p=0.04) *B.pendula* (F_1,4_ =12.3, *p* = 0.025),*Q.robur* (F_1,4_ = 20.9, *p=* 0.010).

**Table 1.**
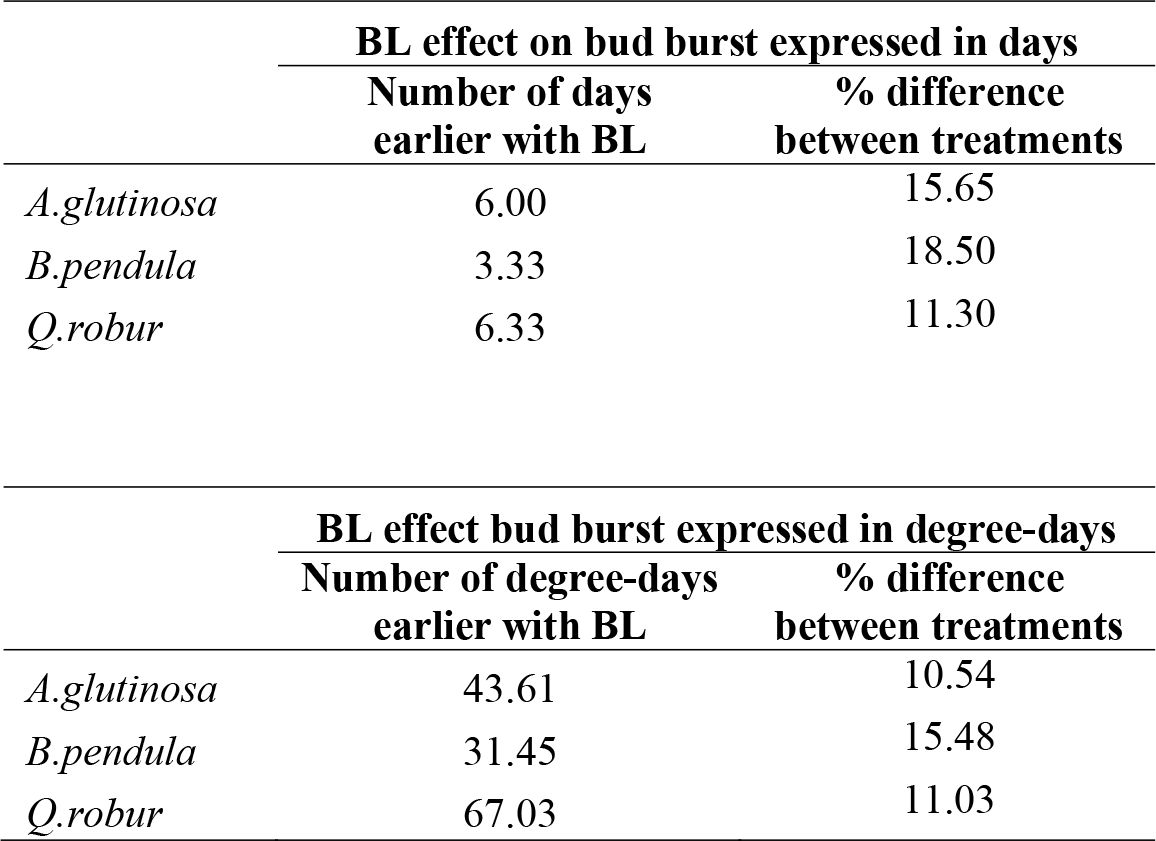
The effect of BL on bud burst expressed both as the difference in the mean number of days and mean degree-days until 50% bud burst in *A.glutinosa, B.pendula* and *Q.robur*, and the percentage differences in the total number of days or degree-day. Degree-days were calculated as the accumulated daily mean temperature > 0°C starting from 1^st^ January 2017.

|  | BL effect on bud burst expressed in days |  |
| --- | --- | --- |
|  | Number of days earlier with BL | % difference between treatments |
| *A.glutinosa* | 6.00 | 15.65 |
| *B.pendula* | 3.33 | 18.50 |
| *Q.robur* | 6.33 | 11.30 |

### What are the mechanisms by which blue light advances bud burst?

Our results demonstrate for the first time that blue light advances spring bud burst in temperate deciduous tree species, through a photoreceptor-dependent mechanism which is yet to be elucidated. This result is consistent with studies of plants of the *Rosacea* family, including *Rosa* sp. and *Prunus cerasifera*, where a higher percentage bud burst and more axillary vegetative shoots and branches were produced under blue light compared to red light (Muleo et al. 2001, Girault et al. 2008). The growth rate and number of preformed leaves in the buds of these species were also higher under blue light. The control of bud burst was exercised in part through the photoregulation of sugar metabolism in this case (Girault et al. 2010), whereby the allocation of sugars to the growing bud increased in the light compared to darkness. The authors attribute the regulation of sugar metabolism in the buds to be primarily mediated by the phytochrome photoreceptors, which have an absorption spectrum in blue light and R:FR, but also possibly by cryptochromes in response to blue light (Reed et al. 1993, Girault et al. 2010). Interestingly, several tree species including *Q.robur, Fagus sylvatica, Betula pubescens, Fraxinus excelsior* and *Acer pseudoplatanus* (Catesson 1964, Barnola et al. 1986, Cottignies 1986, Kelner et al. 1993, Rinne et al. 1994) increase in bud sink strength close to bud burst, resulting in increased mobilization of stem carbohydrates. It appears from these studies that sugar metabolism, at least in part, plays a role in the process of bud burst of tree species.

Regulation of flowering time in response to photoperiod by the blue/UV-A cryptochrome and the R:FR phytochrome photoreceptors has been well established in the model plant *Arabidopsis thaliana* (Guo et al. 1998, Somers et al. 1998). In comparison, much less is known about the role of photoreceptors that regulate bud burst in response to spectral composition. In the tree species *P.tremuloides*, the phytochrome gene PHYB_2_ was found to be coincident with a quantitative trait locus affecting bud-set and bud burst (Frewen et al. 2000). Similarly, cryptochromes have been shown to facilitate blue-light-mediated phenological responses in other plant species (Guo et al. 1998, Meng et al. 2013), so it is reasonable to speculate that cryptochromes and phytochromes could facilitate bud burst in response to blue light. Conversely, although phototropins also absorb blue light, there is no evidence that they mediate phenological responses (Casal 2000). Considering that UV-B can affect bud burst and bud set in *P.tremula* (Strømme et al. 2015), and that plants possess a UV-B-detecting photoreceptor (Rizzini et al. 2011), we suggest that a productive area of future research would be to determine the relative contribution of different photoreceptors to regulating bud burst in response to spectral composition.

### What is the ecological significance of blue light?

We provide two possible explanations (in italics) for the ecological role of blue light advancing bud burst, and why it could have a greater effect on early-successional species, measured in absolute terms as days (or degree days) until bud burst, as opposed to percentage differences. Firstly, it has been demonstrated that blue light enhances photosynthesis in *B.pendula* plantlets *in vitro* (Sæbø et al. 1995), and among other plant species (Goins et al. 1997, Matsuda et al. 2004, Hogewoning et al. 2010). Blue light is often associated with conditions that are favourable to photosynthesis and normal photosynthetic function (Matsuda et al. 2004, Sellin et al. 2011), so it is possible that *blue light advances bud burst in order to allow the plant exploit an opportunity for photosynthetic gains whilst conditions are favourable*. Secondly, the ratio of blue:red light is higher during the twilight hours of sunrise and sunset than the middle of the day (Johnson et al. 1967, Hughes et al. 1984). Robertson et al. (1966) proposes that at higher latitudes the extended hours of twilight rich in blue light and FR could have effects on plant growth not found at lower latitudes, such as the delay in bud dormancy exhibited by *P.abies* grown under day extension using blue light (Mølmann et al. 2006). Given that at high latitudes the day length and thus hours of twilight vary drastically throughout the year, *it is possible that blue light provides cues related to lengthening twilight and daylight periods in spring*. Since photoperiod often has a greater effect on the timing of bud burst in late-successional species (Basler and Körner 2012, Basler and Körner 2014), we might infer that blue light should also have a greater effect on later successional species due to its ecological relevance, i.e. changes in spectral composition related to day length and the length of twilight. Since we used a fixed day length in our experiment, we cannot speculate on the possible interactive effects of blue light and day length here. Similarly, we cannot say whether the bud burst response that we report is related to the proportion of blue light received or the total irradiance of blue light. As such, we suggest future research should test whether bud burst in response to blue light is related to diurnal changes in the proportion of blue light.

Under controlled temperature and light conditions we found that blue light advances bud burst in the tree species *B.pendula, A.glutinosa*, and *Q.robur*. Further research into bud burst and other phenological processes is needed to: establish whether or not the effect of blue light on bud burst is related to diurnal changes in blue light as opposed to a seasonal increase, describe how plants integrate cues from day length and spectral composition (from UV through to far-red) through crosstalk among photoreceptor mediated responses, and combine this information with other environmental cues (e.g. temperature)to produce an environmentally-appropriate response.

## Author Contributions

CCB conceived, designed and carried out the experiment and statistical analyses. TMR supervised the design, analyses and interpretation of results. Both authors wrote the manuscript.

## Acknowledgements

Our acknowledgments to Lammi Biological station staff in particular John Loehr, to Mikko Peltoniemi and the EU Life+ MONIMET phenology camera network, to Viikki Greenhouses Facility at the University of Helsinki and Viikki Arboretum, Marta Pieristé, Marieke Trasser and Ulla Riihimäki for help sampling, Titta Kotilainen for processing the spectral irradiance measurements. Thanks to Otmar Urban and Line Nybakken for giving constructive comments an early version of the manuscript.

